# Behavioural Evaluation of the Anticonvulsant Properties of the Traditional Medicine *Anonna Senegalensis* on Para^bss^ *Drosophila melanogaster Mutants*

**DOI:** 10.1101/519322

**Authors:** Samuel Dare, Emiliano Merlo, Peter E. Ekanem, Jimena Berni

**Affiliations:** Anatomy Unit, Faculty of medicine, Kabale University Uganda; Department of Anatomy, Kampala International University-Western Campus Bushenyi-Uganda; Instituto de Fisiologia y Biofisica Bernardo Houssay, UBA-CONICET, Facultad de Medicina, Universidad de Buenos Aires, Buenos Aires, Argentina; Department of Psychology, University of Cambridge, Cambridge, UK; Anatomy Unit, Institute of Biomedical Sciences, College of Health Sciences, Mekelle University, Mekelle, Ethiopia.; Department of Zoology, University of Cambridge, Cambridge, UK

**Keywords:** *Annona senegalensis*, Bang Sensitive, *Drosophila melanogaster*, epilepsy, *para^bss^*, phenobarbital, phenytoin, seizure

## Abstract

Epilepsy is the most common serious neurological disorder affecting 50 million people worldwide, 40 millions of which live in developing countries. Despite the introduction of a dozen of new Anti-Epileptic Drugs (AEDs), one third of the patients continue to have seizure regardless of receiving a AEDs treatment. This emphasize on the need to discover new drugs with different mechanisms of action. Traditional medicine (TM), pays a significant role in the treatment of epilepsy in many countries and offers an affordable and accessible alternative to AEDs. However, the lack of both empirical testing in animal models and clinical data places constrains to their clinical recommendation.

In this study, we use *Drosophila melanogaster* as a model for epilepsy and tested the anti-seizure effect of leaf and stem bark aqueous extract form *Annona senegalensis*, a plant used as anti-convulsant by rural populations in Africa.

Our results show, that at the concentrations tested, the leaf extract of *A. senegalensis* was more effective than the AEDs phenytoin and phenobarbital to control seizures. These promising results demonstrate that *Drosophila* is an excellent model for new drug discovery and that it could be used to do large scale screening of TMs for the treatment of epilepsy.

## INTRODUCTION

Epilepsy is the most common serious neurological disorder that affects approximately 50 million people worldwide, 40 millions of them living in developing countries.[1,2]. A hallmark of epilepsy are seizures, which are brief episodes of involuntary shaking which may involve a part or the entire body and are sometimes accompanied by loss of consciousness. The episodes are a result of hyper-excitation and/or abnormal synchronization of neuronal circuits’ activity [3]. Seizures can vary in intensity from brief lapses of attention or muscle jerks, to severe and prolonged convulsions. They can also vary in frequency, from less than one per year to several per day.

Recent studies in both developed and developing countries have shown that up to 70% of patients with epilepsy can be successfully treated (i.e. their seizures completely controlled) with anti-epileptic drugs (AEDs) [10]. Phenobarbital is the most commonly prescribed antiepileptic drug in developing countries as recommended by the World Health Organization (WHO) [4,5]. This is probably because phenytoin, carbamazepine, and valproate are up to 5, 15, and 20 times more expensive respectively [6,7]. However, these drugs produce many undesired secondary effects and in 30% of the cases, the available AEDs are ineffective, highlighting the need to develop new treatments [8].

In many developing countries, particularly in Africa and Asia, Traditional Medicine (TM) plays an important role in meeting the demand of primary health care. This is mainly due to its affordability, availability and accessibility. The importance of TM has been recognized by the WHO, which has endorsed an international agreement “the Beijing Declaration” that promises to give TM its place in health systems around the world. The challenge is to “further develop TM based on research and innovation” as stated by one article from the Beijing declaration 2008.

*Annona senegalensis* is commonly found in savannas throughout tropical Africa and its leaves are used by most African rural population as TM for the treatment of epilepsy suggesting that they may have anticonvulsant properties and/or neuroprotective properties. A few studies have tested the activity of AS extracts as anticonvulsant in chemically induced seizures in rodents, supporting a possible role of this TM for the treatment for epilepsy [9-12].

In order to screen for new AEDs it is necessary to use an animal model. In that regard, *Drosophila* adult flies represent a powerful model for the study of seizures [13-16]. This is supported by the similarities between seizures in the fruitfly and humans: (i) all individuals have a seizure threshold; (ii) seizure susceptibility can be modulated by genetic mutations; (iii) the threshold for seizure activity is increased by a previous electroconvulsive shock treatment; (iv) the seizure activity spreads through the central nervous system (CNS) along particular pathways; (v) there is a spatial segregation of seizure activity into particular regions of the CNS; (vi) the seizure phenotypes can be ameliorated by several human AEDs like sodium valproate, phenytoin, gabapentin, and potassium bromide [13].

Furthermore, because there is little redundancy of genes in *Drosophila,* the fruit fly offers a unique opportunity to study human mutated genes in an animal model [17]. It is easy to generate knock-in mutants where the endogenous copy of the gene is replaced by a human version of a mutated gene using homologous recombination or using CRISPR/Cas9 [18]. The new humanised knock-in transgenic flies can then be used for behavioural, physiological or drug testing [17,19].

In this study, we used the *Drosophila* seizure mutant *para*^*bss*^ to investigate the anticonvulsant effect of *A. senegalensis* [20,21] and we compared its effect with two commonly used anticonvulsant phenytoin and phenobarbital.

*para*^*bss*^ is a bang-sensitive mutant, extremely sensitive to seizures that manifest themselves as paralysis and in few individuals it is preceded by as a short period of legs twitching, abdominal muscles contractions, rapid wings flapping and proboscis extensions [22]. In a few animals, paralysis will be interrupted by muscle jerks, in what has been described as a tonic-clonic phase. Within 20 minutes all flies would have recovered and will be capable of maintaining their standing posture and walk or fly.

The seizure phenotype observed in *para*^*bss*^ is due to a gain of function mutation, L1699F, in the sole fly voltage gated sodium channel gene paralytic [22]. This mutation alters the voltage dependence of channel inactivation probably making the neurons more excitable and increasing the risk of seizure [22]. Mutations in the human orthologue, *SCN1A*, are associated with a wide spectrum of epilepsies with over 600 mutations registered in this sole locus [23].

Our results show that *A. senegalensis* leaf extract was more effective to control the seizures than the commonly used AEDs tested. Our empirical testing combined with the one performed in rodents [11-16] are the first step towards the discovery of a new AED that will need further clinical testing. Furthermore, the method presented here is an ideal model for high-thought put screening, which combined with centuries of transmitted knowledge about anti-convulsant TMs, could synergize the discovery of new AEDs for patient irresponsive to the classical treatments.

## RESULTS

### Characterization of the para^bss^ mutant’s phenotype

To evaluating the effect of *A. senegalensis* on seizures, we first characterised the *para*^*bss*^ mutant’s phenotype. To induce seizures, we performed a mechanical shock (10 s vortex) to *para*^*bss*^ and wildtype OrR flies. We evaluated the behaviour of females and males separately because the mutation is located in the × chromosome and sexual dimorphism has been reported [22].

After this stimulus, none of the 60 wildtype flies evaluated showed a seizing phenotype, while 88 ± 5 % of *para*^*bss*^ females and 59 ± 8 % of males were paralysed (t test p= 0.023; Figure 1). After a period seizing, the flies started to recover their standing posture. Finally, all females had recovered at 620 ± 89 s and males at 347 ± 36 s (t test p = 0.029). A mixed ANOVA showed a significant effect of time and an interaction of gender and time. Further *post hoc* analysis showed a significant effect of gender confirming the sexual dimorphism in the seizing phenotype, where males are more resistant to initiate seizures and recover faster than females.

**Figure 1:**
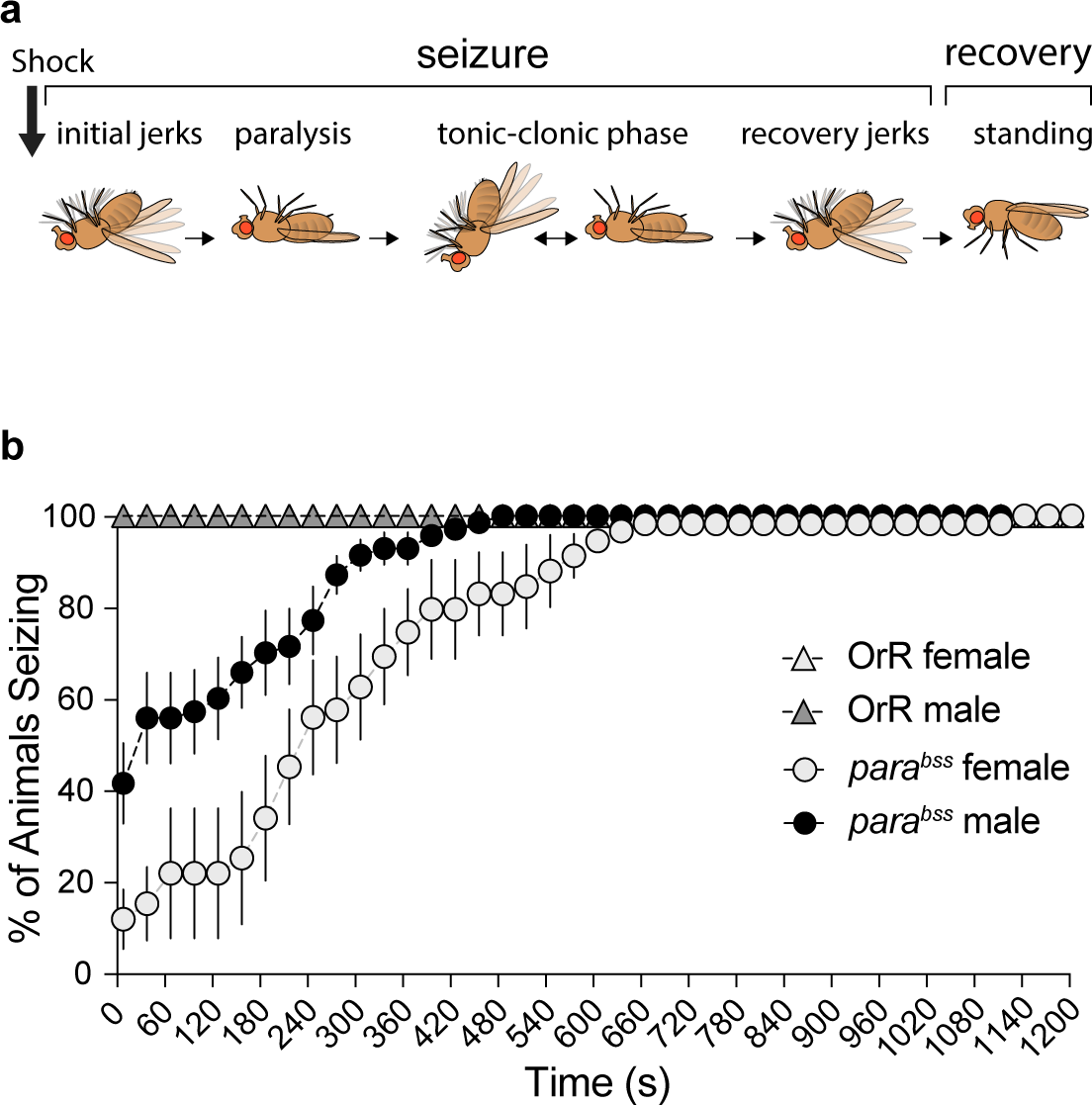
Effect of mechanical shock on seizing behaviour for female and male adult flies of *para*^*bss*^ or OrR genetic background. Mechanical perturbation did not induce seizing in OrR flies. BSS animals showed an immediate increase in seizing that faded as a function of time. Mixed ANOVA on BSS flies behavioural data shows an effect of time (time: F_(2.86, 31.42)_ = 34.03, p < 0.01, η^2^ = 0.76) and an interaction of gender and time (time × gender: F_(2.86, 31.42)_ = 3.17, p = 0.04, η^2^ = 0.22). *Post hoc* analysis shows an effect of gender (F_(1, 11)_ = 11.16, p = 0.007, η^2^ = 0.50). Mean ± SE is shown; *para*^*bss*^ females n = 6; *para*^*bss*^ males n = 7; OrR females n = 3; OrR males n = 3.

This initial data show that the experimental design is particularly well suited to evaluate the dynamics of recovery from seizing and that males and female need to be evaluated separately.

### Annona senegalensis treatment

To evaluate the effect of *A. senegalensis* as anticonvulsant, we decided to compare the behaviour of *para*^*bss*^ flies kept in control food to animals treated with food mixed with aqueous extract from the leaf and stem bark of *A. senegalensis* and two commonly used AEDs, phenytoin and phenobarbital. Flies were left to feed for about 24 hours, transferred to a new empty tube for testing and after a period of acclimation we conducted a seizure experiment as described for Figure 1. We started analysing the behaviour of females. A mixed ANOVA comparing all treatments show a significant effect of time and of the interaction between time and treatment (time: F_(40, 720)_ =75.74, p < 0.001, η^2^ = 0.81; time × treatment: F_(320, 720)_ = 2.25, p, 0.001, η^2^ = 0.50).

We therefore analysed the effect of each drug treatment on the seizure phenotype. As expected, when *para*^*bss*^ flies were kept in control food, with no drugs, the mechanical stimulation induced seizures in a high percentage of flies, most of which recover during the first 600s with only few flies seizing for a longer period of time.

As a positive control we first evaluated phenytoin, a drug known to modulate the gating of voltage gated sodium channels [24] and that has already been shown to diminish seizure on *para*^*bss*^ flies, but which had been tested in different conditions [25]. Comparing the control with two concentrations of Phenytoin showed a significant effect of time and an interaction of time and treatment (time: F_(40, 240)_ = 17.00, p < 0.001, η^2^ = 0.74; time × treatment: F_(80, 240)_ = 4.58, p < 0.001, η^2^ = 0.60). Follow on analysis were not significant when comparing between treatments only. The lack of significance of the treatment might be due to the high variability of the data and the fact that the higher concentration of phenytoin seems to slightly worsen the recovery from seizure. However, the analysis of the percentage of flies seizing at t = 0 showed that the drug was different from the control (ANOVA F_(2, 6)_ =8.53, p < 0.018, η^2^= 0.74) having a protective effect on the initiation of seizures at both concentrations (control to phenytoin at 1.7 mg.ml^-1^ p = 0.035; at 3.4 mg.ml^-1^ p = 0.022). And this initial difference is maintained over time with significantly fewer flies seizing compare to control for each time point from t = 0 until t = 210s for 4.76 mg.ml^-1^ and until t = 270 s for 9.09 mg.ml^-1^ (*post hoc* Tukey analysis, figure 2A).

**Figure 2:**
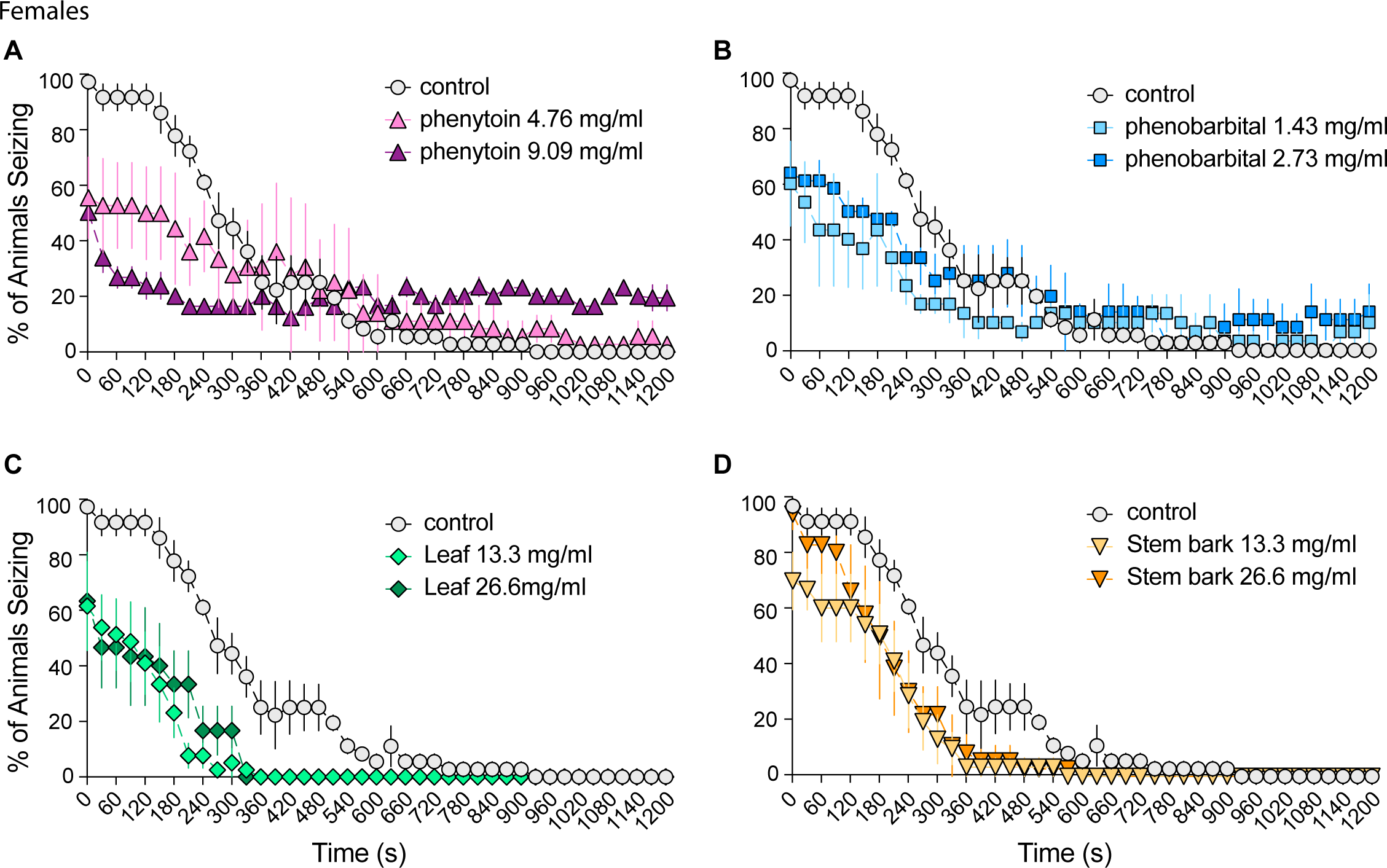
Effect of phenytoin, phenobarbital and A. senegalensis extracts on seizures of *para*^*bss*^ females. In *para*^*bss*^ flies treated with the control food, mechanical perturbation induced seizing that decreased over time. Phenytoin and phenobarbital treatment at both concentrations decreased the percentage of seizing flies at the beginning of the experiment but the time of recovery was not improved. *A. senegalensis* leaf extract decreased the percentage of flies seizing at the beginning of the experiment and also decreased the time to recovery. The treatment with stem bark extract had a weaker effect but the percentage of flies seizing was below the curve of the control treatment at many time points. A mixed ANOVA was performed to compare all treatments, because it was significant follow on mixed or single way ANOVAs were performed comparing specific treatments or time. Tuckey *post hoc* analysis for each time point was performed. See text for values. Mean ± SE is shown; n = 3 for all treatments.

To test if drugs acting on different targets are also capable of improving the seizure phenotype of *para*^*bss*^ flies, we then tested the anti-convulsant drug phenobarbital that mainly acts via modulation of GABA_A_ receptors [24]. When comparing *para*^*bss*^ flies in control food with flies treated with two concentrations of phenobarbital, the decay of the percentage of seizing animals over time was found significant (time: F_(40, 240)_ = 43.24, p < 0.001, η^2^ = 0.88) for the three manipulations and an interaction between time and treatment was also significant (time × treatment: F_(80, 240)_ = 3.67, p < 0.001, η^2^ = 0.55). Further *post hoc* Tukey analysis demonstrated the effect of phenobarbital on *para*^*bss*^ flies, which is significant at from t = 0 until t = 250 s for 1.43 mg.ml^-1^ and t = 210 s for 2.73 mg.ml^-1^ of phenobarbital (figure 2B). After this initial period, a high proportion of the flies have recovered and no differences between the treatment are observed. This similarity over most of the experimental time is confirmed by the non-significance of a follow-on ANOVA when the controls are compared to the treatments independently of time.

We then tested the effect of the leaf extract of *A. senegalensis* (Figure 2C). As per the other treatments, a significant effect of time and an interaction of time and treatment was shown (time: F_(40, 240)_ = 49.56, p < 0.001, η^2^ = 0.89; time × treatment: F_(80, 240)_ = 4.02, p < 0.001, η^2^= 0.57). A follow-on ANOVA also showed a significant effect of treatment (F_(2,6)_ = 17.83, p = 0.003, η^2^ = 0.86) and *post hoc* Tukey analysis showed that the control was significantly different from both concentrations of the leaf treatment (comparison control to leaf 13.3 mg.ml^-1^: p = 0.004; control to leaf 26.6 mg.ml^-1^: p = 0.006). The two leaf concentrations did not produce a different effect on seizure between each other. This result supports an anti-seizure effect of the leaf extract of *A. senegalensis*, which is more robust than the one observed with the classical AEDs. The anti-seizure effect of *A.* senegalensis can be divided in two related aspect that are ameliorated. The first on is the initiation of seizure at t = 0, which is significantly decreased in treated flies for both concentrations (*post hoc* Tukey analysis control to leaf p < 0.001 for both concentrations). This is followed by a period of recovery where a higher percentage of controls seize compare to leaf treated flies until t = 480 s for both concentrations. In this period, the rate of recovery does not seem to be improved by the leaf treatment, but because the starting point was lower, *A. senegalensis* shortens the time to complete recovery (% seizing = 0) (F = 18.59, p =0.003, R^2^ = 0.86) for both concentrations (comparison control to leaf 13.3 mg.ml^-1^: p = 0.004; control to leaf 26.6 mg.ml^-1^: p = 0.005).

Finally, we evaluated if an extract from the stem bark of *A. senegalensis* would produce the same effect than the leaf extract. This was not the case, because there was a significant effect of time (time: F_(40, 240)_ =72.04, p < 0.001, η^2^ = 0.92) and interaction between time and treatment (time × treatment: F_(80, 240)_ = 1.54, p = 0.007, η^2^ = 0.34) but the effect was not strong enough to be independent of time. Despite the similarities between the three curves, the stem bark treated percentage of seizing flies is consistently below the one of controls for the first half of the experiment and this is reflected by significant differences when performing a *post hoc* Tukey analysis (control compared to 13.3 mg.ml^-1^ of stem bark extract, p<0.05 for all times from t = 0 until t = 480s; control compared to 26.6 mg.ml^-1^ p<0.05 from t = 120 s until t = 360s; Figure 2D). However, these differences are smaller than in the case of the leaf treatment showing a much weaker effect.

We then evaluated the seizure phenotypes of males exposed to the same treatments than the females. In males, the percentage of flies having seizure after the mechanical shock is significantly lower than in females (figure 1) and at around 240 seconds most of the fleis have recovered from seizure. Therefore, significant effects due to the treatment were expected to be found at the beginning of the experiment. A significant effect of time and of the interaction between time and treatment was found (time: F_(40, 760)_ =17.00, p < 0.001, η^2^ = 0.47; time × treatment: F_(320, 760)_ = 1.91, p < 0.001, η^2^ = 0.45).

We therefore proceeded to perform statistics for each drug. When comparing the control group to flies treated with two concentrations of phenytoin a significant effect of time and of the interaction between time and treatment were found (time: F_(40, 280)_ =9.15, p < 0.001, η^2^ = 0.57; time × treatment: F_(80, 280)_ = 3.40, p, 0.001, η^2^ = 0.49), but follow on analysis comparing treatment was not significant. A significant difference (p< 0.05) was found at the beginning of the experiment from t = 0 until t = 120 for 4.76 mg.ml^-1^ and until t = 90 s for 9.09 mg.ml^-1^ (*post hoc* Tukey analysis, figure 3A).

**Figure 3:**
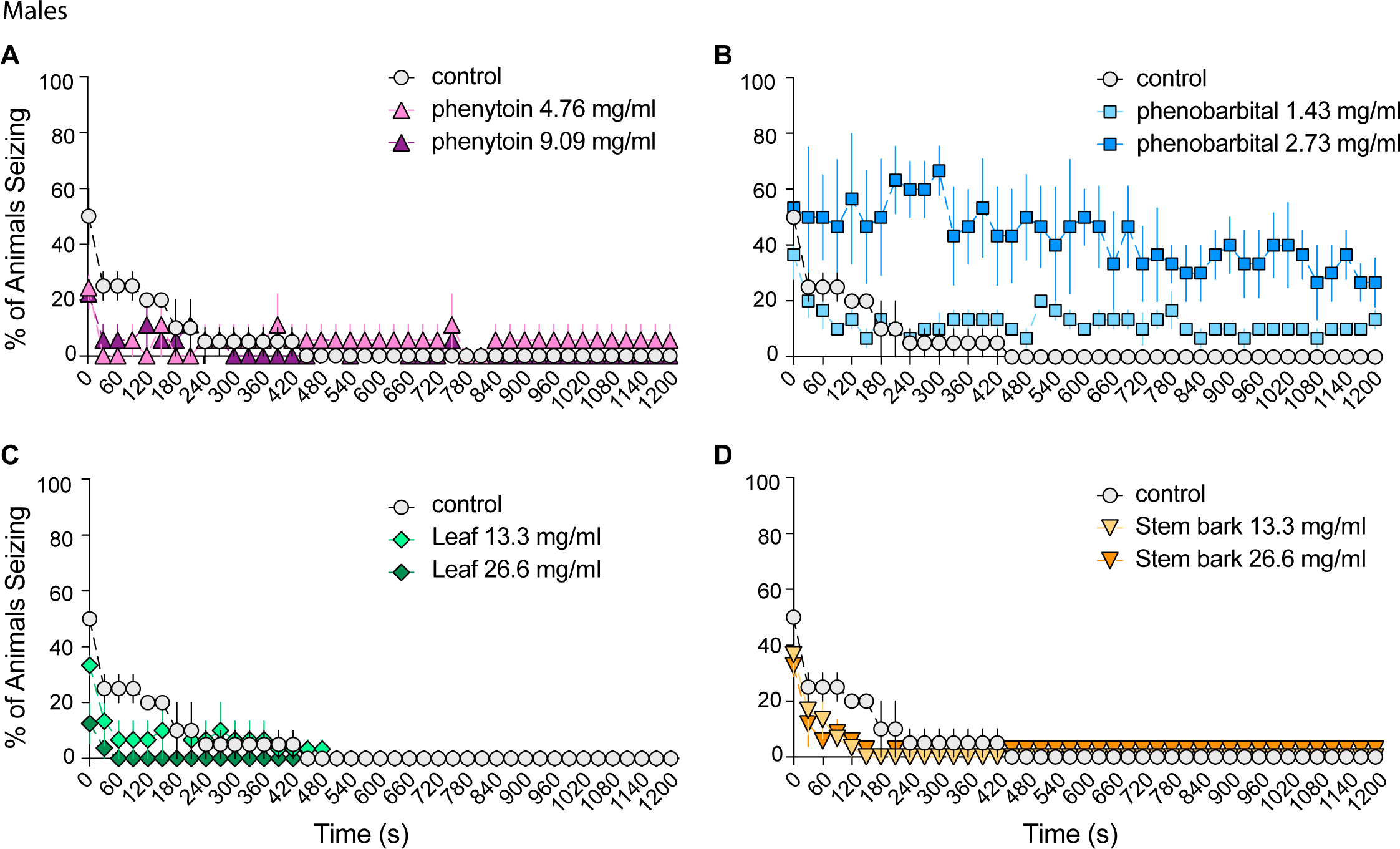
Effect of phenytoin, phenobarbital and *A. senegalensis* extracts on seizures of *para*^*bss*^ males. As expected, mechanical chock on *para*^*bss*^ male flies induced a seizure phenotype that was less penetrant than in females. Phenytoin protected the flies from seizure and the animals recovered very fast. Phenobarbital did not show an effect at the lower concentration and worsened the seizures at the highest concentration. The leaf extract improved the seizures at the beginning of the experiment as did the stem bark extract. A mixed ANOVA was performed to compare all treatments, because it was significant follow on mixed or single way ANOVAs were performed comparing specific treatments. Tuckey *post hoc* comparison for each time point was performed. See text for values. Mean ± SE is shown; n = 2 for *para*^*bss*^ control, n = 3 for all the other treatments.

With phenobarbital, the percentage of seizure significantly changed over time (time: F_(40, 80)_ = 7.11, p < 0.001, η^2^ = 0.50) and there was an interaction between time and treatment (time × treatment: F_(80, 280)_ = 2.25, p < 0.001, η^2^ = 0.39). This difference could be attributed to treatment effects (F_(2, 7)_ = 8.29, p = 0.014, η^2^ = 0.70) and in particular to a deterioration of the flies health compare to controls at the higher concentration of phenobarbital where seizures proceed until the end of the experiment (control to phenobarbital 10mg.ml^-1^ p = 0.014).

When evaluating the effects of the treatments with leaf extract from *A. senegalensis,* we found that the percentage of seizure of the three groups, control, leaf 13.3 mg.ml^-1^ and leaf 26.6 mg.ml^-1^, changed over time (time: F_(40, 280)_ =12.63, p < 0.001, η^2^ = 0.64) and there was a treatment effect over time (time × treatment: F_(80, 280)_ =2.80, p < 0.001, η^2^ = 0.45). *As per* phenytoin, there was a significant difference at the beginning of the experiment between the percentage of seizing flies in the control and in the leaf treatments (control compared to leaf extract 13.3 mg.ml^-1^ p<0.001 at t = 30 s, 60 s and 90 s; compared to leaf extract 26.6 mg.ml^-1^ p<0.001 from t = 0 s until t = 150 s).

A similar effect of time and interaction of treatment over time was also present with the stem bark of *A. senegalensis* but the effect were not sufficiently large to be significant independently of the time (time: F_(40, 280)_ = 21.33, p < 0.001, η^2^ = 0.75; time × treatment: F_(80, 280)_ = 2.62, p < 0.001, η^2^ = 0.43). However, there was a significant improvement of the seizing phenotype at the beginning of the experiment with the stem bark treatment at 13.3 mg.ml-1 been significantly different (p< 0.05) from t = 0 s until t = 210 s, excluding t = 30s and at higher concentration, the differences spanned from t = 0 until t = 180 s.

Taken together these results show that both in females and males, there was an effect of treatments over time for the commonly used AEDs as well as for the stem bark and leaf extract from *A. senegalensis*. The leaf treatment was the strongest for both sexes as reflected by its ability to very significantly protect against seizure and to also improve the recovery time in females.

## Discussion

Up to date, approximately 30 AEDs are approved to treat patients with a disease of very complex and variable aetiology. However successful to control seizures in most of the patients, a non-negligible percentage of them experience undesired secondary effects and 30 percent of patients will still present seizures, highlighting the necessity of discovering new drugs.

Almost all the commonly used AEDs were discovered by screening in animal models and their mechanisms of action were also studied in non-human models, emphasising the importance of animal testing for drug discovery in epilepsy. Over the years, medicinal plants have been used for treatment of various diseases including epilepsy. The knowledge gathered and transmitted for generations about the anti-seizure properties of herbs is a very rich source of information for new drugs discovery. However, an empirical testing of their efficacy and toxicity is required. A few hundred drugs from TM have been tested in animal models but the lack of follow up clinical studies places constraints on the clinical recommendation of herbal medicine [26].

In the present study, we have used a *Drosophila* seizure model to test the effect of *A. senegalensis* as an anti-seizure drug [13]. We decided to use a fly with a gain of function mutation in the unique voltage gated sodium channel of *Drosophila* genome, *para*^*bss*^, which is extremely sensitive to seizures. This mutation is generally accepted as a model for intractable epilepsy due to the complexity of the seizures, including a tonic-clonic-like phase, and because it is largely resistant to treatment with AEDs [22].

In our hands, *para*^*bss*^ showed the expected seizure phenotype and we observed a higher percentage of females seizing compare to males. The sexual dimorphism could be due to differences in the level of expression of *para*^*bss*^ (located in the × chromosome, female have two copies, males one) but because there are dose compensation mechanisms [27], we favour the idea that it is due to endocrine differences. In humans, sexual differences in seizure type and symptoms are well known [28] and certain types of epilepsy are gender-specific like catamenial epilepsy with seizures clustered around the perimenstrual and periovulatory period. Animal models should be used more widely in the investigation of such sexual differences and *Drosophila* could also play its part.

We therefore continue our experiments analysing separately males and females. Our experimental design was particularly suited to analyse two main effects of the treatment. First, the protective effect, where the number of flies seizing after the mechanical shock could diminish compare to controls, and second, the recovery effect, as the time to recover standing posture could be shorten. To quantify the effect of *A. senegalensis*, we decided to compare it to two commonly used AEDs. As a positive control, we chose phenytoin because it effects had already been studied, but at different concentrations and in populations where males and females were mixed together [25]. Phenytoin specifically targets voltage gated sodium channels inducing a non-conductance state of the channel similar to inactivation [24]. We also tested phenobarbital, which had not been tested before on *para*^*bss*^ flies and acts on GABA_A_ and glutamate receptors and HVA calcium channels. Phenobarbital main anti-seizure effects is due to a positive allosteric modulation of GABA_A_ receptors, shifting the relative proportion of open states, favouring the longest live open state associated with prolonged bursting [24].

In flies treated with the commonly used AEDs, we observed the expected change of symptom, being phenytoin the most effective. A clear protective effect with fewer flies seizing compare to controls at the beginning of the experiment t = 0 was quantified. This was significant for phenytoin and phenobarbital for both concentrations and sexes apart for phenobarbital at the higher concentration that seems to have produced a toxic effect on male flies.

The treatment with Leaf extract of *A. senegalensis* was the most effective treatment we tried on seizures. Both concentrations showed a significant protective effect decreasing the percentage of flies seizing at t = 0 by more than 35 % for both concentrations. It also was the only drug showing a significant shortening of the time to recovery by 60 % for both concentrations. In males, we also quantified the protective effect of leaf extract of *A. senegalensis* but not the recovery time. An experimental design where recovery time could be prolonged for *para*^*bss*^ flies would be required to confidently test the effect of *A. senegalensis* on that parameter.

In comparison, the treatment with the stem bark extracts was less effective suggesting the active compound is less concentrated in the stem bark than in the leaves.

Several studies have evaluated the effect of *A. senegalensis* extracts prepared with different solvents and from several parts of the plants (leaves and root) on rodents. They have shown protective effects in pentylnetetrazole and maximal electroshock induced seizure but no further mechanism has been studied [11-16].

Taken together, these results are very promising and are laying the bases for future studies aiming to discover the active compound with anti-seizure effect that is present in the leaf extract of *A. senegalensis.* The simplicity, reproducibility and low cost of the testing method makes it ideal to test the effect of sub-extracts of the leaf until finding the fraction where the active compound is present. This could then be purified and analysed with mass-spectroscopy. In that direction, a possible candidate that should be tested is Kar-16a-19oic acid, a diterpenoid with sedative and anti-seizure properties [9] that has been purified from methanol-methylene chloride extracts from the root bark of *A. senegalensis.* This compound has a very low toxicity (Kar-16a-19oic acid LD_50_ = 3800 mg/kg compare to LD_50_ = 1635 mg/kg for phenytoin and LD_50_ = 660 mg/kg for phenobarbital (oral administration, Merk index)), which could make it ideal for its clinical recommendation as a TM [9]. Thanks to the high structural and functional conservation that exist between channels and more generally proteins found in *Drosophila* and humans, once the anti-seizure compound found, *Drosophila* offer and array of possibilities for the investigation of the mechanisms of action of the drug (reviewed by [14])

In conclusion, our work shows how powerful *Drosophila* is as a model for screening of AEDs. Regardless the fact that there was little information about how to prepare the plant extract, we were able to find a strong anti-seizure effect of the leaf extract from *A. senegalensis*. This method could be used for high-through put screening of many TM used as anti-convulsion treatment giving hope to the patients with intractable epilepsy.

## MATERIALS AND METHODS

### Indentification and Extraction of Annona Senegalensis

*Annona Senegalensis* leaves and stem barks were collected from the Northern part of Uganda and were identified at Biology department of Mbarara University of Science and Technology Uganda. Extraction was performed at the Pharmacology department laboratory of School of Health Sciences, Kampala International University Western Campus, Ishaka Uganda. Specifically, dry leaves and stem bark of plant were grounded using a blender to obtain 200 g powder which was mixed with 1000 ml distilled water in a sterile conical flask. The mixture was placed on a shaker for 48 hours after which it was sieved to remove the solid particles. The remaining liquid was filtered using Wattman no. 1 filter paper. The filtrate was placed in a water bath at 35 ^o^C for 1 week to evaporate excess water and to obtain a dry powder [29]. The extract obtained was stored in refrigerator until the experiments was performed.

### Flies Treatment and Evaluation

Flies of the bang sensitive family, in particular *bang senseless* (*bss*) mutants of the paralytic gene *para*^*bss*^, obtained from Richard Baines laboratory, were used for this study [13,14]. Young flies were anesthetized by cold exposure in a freezer (−4°C) until anesthetized, separated into males and females with 9-12 per vial on a cold plate. They were then transferred into vials with standard cornmeal food with or without drugs. Flies were allowed to feed for approximately 24 hours prior to the experiment in their respective vials. The day of the experiment, the flies were transferred into empty vials, allowed to rest for 15 to 30 minutes and then mechanically stimulated by placement on a bench-top vortex at maximum speed for 10 seconds. The number of flies on their backs; paralyzed, shaking or standing were recorded at 30 seconds’ intervals for 20 minutes. Paralysed + shaking flies is considered as seizing and full recovery of flies was defined as standing. Experiments were repeated 3 times apart from for the characterization experiment where *para*^*bss*^ female n = 6 and male n = 7 and in male control food n = 2.

### Drug preparation for fly administration

Drugs were administered mixed with standard cornmeal food. The concentration used was calculated from the amount of drug mixed in 5 ml of fly food. For *A.Senegalensis,* 80 mg and 160 mg of powder extract was dissolved in 1 ml distilled water, which was mixed with 5 ml of fly food for 13.33 and 26.66 mg/ml respectively. For the drugs, one tablet of 30 mg of Phenobarbital (B-Tone; Kampala Pharmaceuticals Industries (1996) Ltd.) or one tablet of 100 mg of Tophen Phenytoin; AgogPharma Ltd., India) was dissolved in 1 ml of water and 250 ul or 500 ul of drug was then mixed in 5ml of fly food for the low and high concentrations respectively.

### Statistical analysis

All data was expressed as the mean ± SEM. T test or One-way ANOVA were used to compare % seizing at t = 0 and time for 100% recovery. Repeated-measures ANOVAs, with “Group” as the between-treatment factor and time as the within-subjects factor. Tukey’s *post hoc* test comparisons were used for further analysis. Sphericity was considered in all data set [33].

## Author Contributions

S.D and J. B conceived and designed the experiments, S.D. performed the experiments, E.M did the data analysis and statistics, P.E.K. prepared the extract; J.B. wrote the manuscript with contributions from all authors.

## ACKNOWLEDGEMENT

We thank Nara Muraro for constructive comments on the manuscript and Richard Baines for sharing the *para*^*bss*^ flies. Nan Hu for his help with the *para*^*bss*^ control experiment. We thank the Institute of Biomedical Research Laboratory, Kampala International University (KIU) Western Campus Uganda and Mbarara University of Science and Technology for providing laboratory space and Trend in Africa for setting up the laboratory and for organising the first courses of neurogenetics in *Drosophila* in Africa where S.D. was trained. We thank the Cambridge in Africa programme the Alborada Trust Fund for funding the project. E.M. is funded by CONICET, Argentina; J.B. is funded by a Sir Henry Dale Fellowship (Wellcome Trust and the Royal Society) Grant 105568/Z/14/Z.

